# The design and analysis of binary variable traits in common garden genetic experiments of highly fecund species to assess heritability

**DOI:** 10.1101/018044

**Authors:** Sarah W. Davies, Samuel V. Scarpino, Thanapat Pongwarin, James Scott, Mikhail V. Matz

## Abstract

Many biologically important traits are binomially distributed, with their key phenotypes being presence or absence. Despite their prevalence, estimating the heritability of binomial traits presents both experimental and statistical challenges. Here we develop both an empirical and computational methodology for estimating the narrow-sense heritability of binary traits for highly fecund species. Our experimental approach controls for undesirable culturing effects, while minimizing culture numbers, increasing feasibility in the field. Our statistical approach accounts for known issues with model-selection by using a permutation test to calculate significance values and includes both fitting and power calculation methods. We illustrate our methodology by estimating the narrow-sense heritability for larval settlement, a key life-history trait, in the reef-building coral *Orbicella faveolata*. The experimental, statistical and computational methods, along with all of the data from this study, were deployed in the R package multiDimBio.

## Introduction

Organisms with high fecundity, small propagule size, and limited parental investment, also referred to as r-selected species, often exhibit higher levels of nucleotide diversity and/or standing genetic variation when compared to k-selected species (Romiguier *et al.*, 2014). Many marine species, including fish and invertebrates, exhibit these r-selected life history characteristics (Doherty & Fowler, 1994) and indeed have been shown to exhibit high levels of genetic diversity (Bay *et al.*, 2004; Davies *et al.*, 2015). However, this high genetic diversity does little to predict how a population will respond to environmental perturbations, such as those caused by climate change. Instead, the key question is not how much variation is present, but what is the heritability of the traits under selection following the perturbation. More specifically, we focus on narrow-sense heritability, defined as the proportion of phenotypic variance attributable to additive genetic effects (Lynch & Walsh, 1998).

Quantifying narrow-sense heritability for non-model organisms presents both experimental and statistical challenges. Most experiments aiming to quantify narrow-sense heritability involve multi-generational breeding programs and large numbers of crosses with many culture replicates to account for “jar effects,” both of which are rarely feasible for non-model marine species. Here we present a quantitative genetic methodology for binary traits in highly fecund species. The method does not require the onerous sampling schemes usually required for these types of experiments. Instead, our approach leverages high fecundity to perform a single bulk multi-sire cross, where larvae are sorted according to the binary trait of interest. Single larvae that “succeeded” and “failed” are then genotyped and their sire assignments are compared to the predicted distribution of sire assignments in the original design. The narrow-sense heritability of these data can be estimated using a generalized linear mixed model with a binomial error distribution. However, as we discuss below, appropriately determining statistical significance is non-trivial. This method of quantifying heritability of binary traits is broadly applicable to many traits of interest including–but not limited to–stress tolerance, dispersal potential, and disease susceptibility.

Here, we show the application of this method to investigate the heritability of dispersal potential in reef-building coral larvae. The majority of corals, like many other marine invertebrates, release gametes into the water annually that develop into planktonic larvae that are dispersed by ocean currents, representing the coral’s only dispersal opportunity (Baird *et al.*, 2009). These pelagic larvae can travel great distances before settling on a reef but once the larva settles in a location, it will remain there for the duration of its life. Selection for dispersal potential is therefore limited to optimizing larval traits, which can be investigated through classical quantitative genetics, e.g. Meyer *et al.* (2009). With this new method, we aimed to estimate how much variation in the early larval responsiveness to settlement cue depends on the genetic background of larvae. The experiments were performed on larvae of the hermaphroditic lesser star coral, *Orbicella faveolata*, which is an important Caribbean reef-building coral. To analyze these data, and specifically estimate the narrow-sense heritability of this binary trait, we developed a Monte Carlo method for performing model selection and calculating statistical power with generalized linear mixed models. The code and data are available in the R package multiDimBio (Scarpino *et al.*, 2014).

## Materials and methods

### Experimental Methods

#### Crossing design and larval rearing

One day prior to the annual coral spawn on August 7, 2012, ten independent *O. faveolata* colony fragments (10cm × 10cm) were collected from the East Flower Garden Banks National Marine Sanctuary, Gulf of Mexico. Colonies were maintained in flow through conditions aboard the vessel shaded from direct sunlight. Colonies were at least 10m apart to avoid sampling clones. Prior to spawning, at 20:00CDT on August 8, 2012, colonies were isolated in individual bins filled with 1*μ*m filtered seawater and shaded completely. Nine colonies spawned at approximately 23:30CDT and gamete bundles were collected and eggs and sperm were separated with nylon mesh. Only sperm from these colonies were used in the crossing design where each served as an independent sire. Samples from each sire were preserved in ethanol for genotyping. Divers collected gamete bundles directly from three colonies during spawning and eggs were separated and served as maternal material (N=3 dams). Eggs were divided equally amongst fertilization bins (N=9 per dam) and sperm from each sire was added at 0200CDT on August 9, 2012 for a total of 27 fertilization bins. Control self-cross trials verified that self-fertilization was not detectable in our samples. After fertilization, at 0800CDT, excess sperm was removed by rinsing with nylon mesh, and embryos for each dam across all sires were pooled in one common culture. Densities were determined and larvae were stocked into three replicate culture vessels at 1 larva per 2ml for a total of nine culture containers (N=3 per dam). Larvae were transferred to the University of Texas at Austin on August 10, 2012. Following spawning, colony fragments were returned to the reef. All work was completed under the Flower garden Banks National Marine Sanctuary permit #FGBNMS-2012-002.

#### Common Garden Settlement Assay

On August 14, 2012, 6 day old, pre-competent larvae from three replicate bins for one dam were divided across three settlement assays. 400 healthy larvae per culture replicate were added to a sterile 800ml container with five conditioned glass slides and finely ground, locally collected crustose coralline algae (CCA), a known settlement inducer for this coral genus (Davies *et al.*, 2014). Cultures were maintained for three days after which each slide was removed and recruits were individually preserved in 96% ethanol, representing larvae exhibiting “early” responsiveness to settlement cue. Culture water was changed, new slides were added with additional CCA and larvae were maintained until they reached 14 days old. All settled larvae on slides were discarded and 50 larvae per culture were individually preserved in 96% ethanol. Larvae from the other two dams were not used in these assays due to high culture mortality.

#### Larval DNA Extraction

Larval DNA extraction followed a standard phenol-chloroform iso-amyl alcohol extraction protocol, see Davies *et al.* (2013), with modifications to accommodate for the single larva instead of bulk adult tissue.

#### Parental Genotyping

Sire genotyping was completed using nine loci from Davies *et al.* (2014) and four loci from Severance *et al.* (2004) following published protocols. Amplicons were resolved on agarose gel to verify amplification and molecular weights were analyzed using the ABI 3130XL capillary sequencer. GeneMarker V2.4.0 (Soft Genetics) assessed genotypes and loci representing the highest allelic diversities amongst the sires were chosen as larval parentage markers.

#### Larval Parentage

To compensate for the low larval DNA concentrations, 3*μ*l of each single extracted larva (unknown concentration) was amplified in a multiplex reaction with six loci from Davies *et al.* (2013) with the following modifications: 1*μ*M of each fluorescent primer pair (N=6) and 20*μ*L reaction volumes (Table 1). Alleles were called in GeneMarker V2.4.0 and offspring parentage was assigned based on presence/absence of sire alleles. Data were formatted into a dataframe consisting of the number of early settlers and swimming larvae that were assigned to each sire (A-J) from each of three replicate bins (1-3).

**Table 1.**
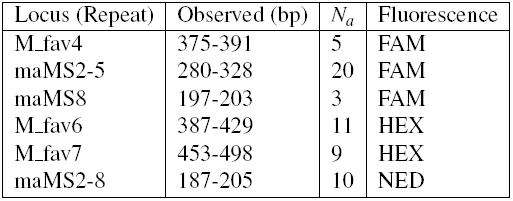
Summary of microsatellite used in paternity assignment

| Locus (Repeat) | Observed (bp) | $N_a$ | Fluorescence |
| --- | --- | --- | --- |
| M_fav4 | 375-391 | 5 | FAM |
| maMS2-5 | 280-328 | 20 | FAM |
| maMS8 | 197-203 | 3 | FAM |
| M_fav6 | 387-429 | 11 | HEX |
| M_fav7 | 453-498 | 9 | HEX |
| maMS2-8 | 187-205 | 10 | NED |

### Statistical Methods

#### Estimating narrow-sense heritability from binary data

In principle, estimating narrow-sense heritability for a binomially distributed trait, such as coral settlement, is straightforward, see Gilmour *et al.* (1985); Biscarini *et al.* (2014, 2015). The desired quantity is the among-sire variance, denoted as *τ*^2^, which can be estimated using a generalized linear mixed model with a binomial error distribution.

Suppose we have binary observations *y*_*ij*_ ϵ{0, 1} where *i* index units (sires) and *j* indexes observations within units. The model is simple Bernoulli sampling, parametrized by log odds:

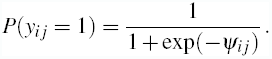

We will assume that the log odds have a sire-level random effect:

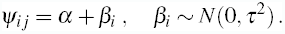

Thus we have a simple binary logit model with a single random effect. A standard result on logit models is that we can represent the outcomes *y*_*ij*_ as thresholded versions of an latent continuous quantity *z*_*ij*_ (Holmes *et al.*, 2006):

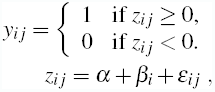

where *ε_i j_* follows a standard logistic distribution. Thus we can interpret narrow-sense heritability *θ* as the ratio

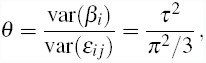

exploiting the fact that the variance of the standard logistic distribution is *π*^2^*/*3. This provides a natural interpretation of heritability of a binary trait on a scale (log odds) familiar from ordinary logistic regression. As a result, the among-sire variance can be transformed into narrow-sense heritability by dividing *τ*^2^ by 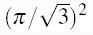 and then multiplying that quantity by two.

However, under this model, determining whether statistical support exists for an among-sire variance greater than zero remains a challenge. Traditionally, an approach to the problem would be to fit two models, one where *τ*^2^, the among-sire variance, is a free parameter and one where it is constrained to zero. These models can then be compared, and model selection performed, using a likelihood ratio test, or in this case the difference in each model’s deviance, which is equivalent to a likelihood ratio test for nested models. When computing a p value for such a likelihood ratio test, the difference in deviance is assumed to follow a chi-squared distribution with degrees of freedom equal to the difference in the number of free parameters. Unfortunately, this assumption is invalid when the null hypothesis exists at the boundary of possible parameter values (Self & Liang, 1987), as is the case with narrow-sense heritability. Instead, our approach is to construct a permutation-based method for calculating a p value for the likelihood ratio test and performing model selection.

#### Monte Carlo simulation for the likelihood ratio test

Briefly, we fix the probability of settlement, *p*_*settle*_, to be equal across all sires, in this case *p*_*settle*_ = 0.285 (the global mean), and simulate 1,000 data sets where the number of offspring for each sire in each of three bins is drawn from a negative binomial distribution with *μ* = 4.63 and 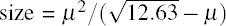 again these are the empirically observed values across sires. The resulting 1,000 data sets have the same structure as the observed data, but the only among sire variability comes from sampling, the true *τ*^2^ = 0. For each simulated data set, we fit both the null and alternative models, described above, and calculate the likelihood ratio, difference in model deviances. This null distribution of differences can then be used to calculate a p value by asking how many of these simulated values are greater than the observed values. Importantly, while computationally intensive, this approach does not suffer from the problem of using the aforementioned asymptotic approximation to the chi-squared distribution.

#### Power analysis

With any novel experimental design, it is desirable to construct a method for estimating its statistical power. Using the Monte Carlo approach designed to calculate p values for likelihood ratio tests, we can simulate data sets with an arbitrary number of sires, number and variance in offspring, among-sires variance, and number of bins. By repeatedly simulating data sets using fixed combinations of these parameters, the statistical power is simply the fraction of times we correctly reject the null hypothesis. Similarly, the false positive rate is the fraction of times we falsely reject the null hypothesis.

#### Implementation

All code and data developed for this study are available in the R package multiDimBio (Scarpino *et al.*, 2014).

## Results

### Parentage

Larvae that amplified for>2 loci were considered successful amplifications. A total number of 55 recruits (binary successes) were collected and of these 47 were amplified and 37 were assigned parentage. A total number of 129 swimming larvae (binary failures) were extracted and of these 112 amplified successfully and 81 were assigned parentage.

### Statistics

Using the described experimental design and statistical methods, we were unable to detect a significant random effect of sire, although there was a trend in overall variation in early settlement among sires (Figure 1). However, considering the number of sires used and offspring sampled in our study, the actual narrow-sense heritability would have to be greater than 0.6 to achieve 80% power (Figure 2a). However, this experimental set up should be sufficiently powered to correctly fail to reject the null hypothesis if in fact true among sire variance was zero (Figure 2b).

**Figure 1.**
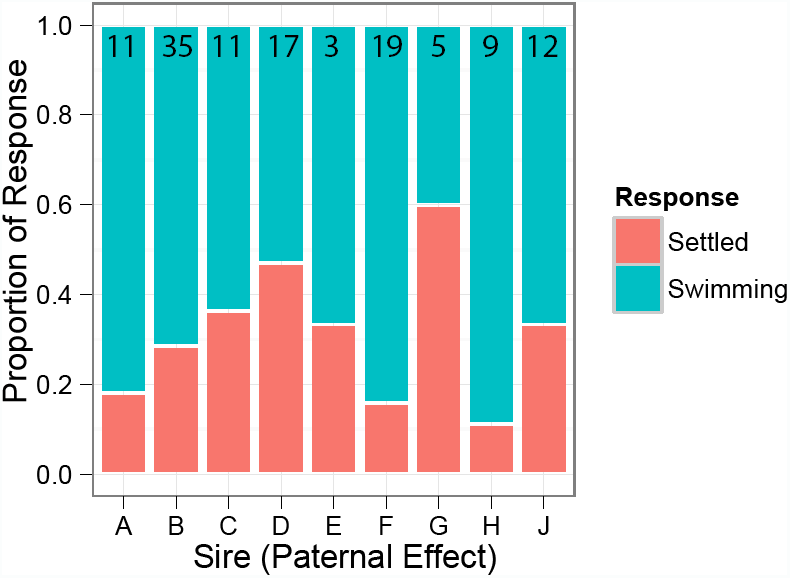
Proportion of settled (successes) and swimming (failures) larvae belonging to each sire. The total number of genotyped larvae assigning to each sire is indicated at the top of each bar.

**Figure 2.**
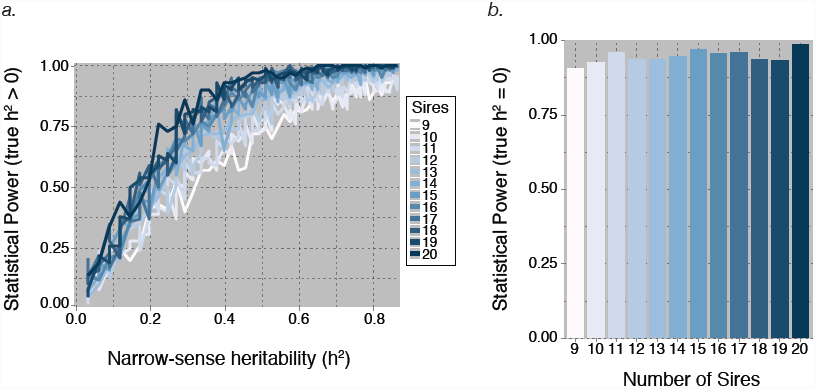
Power analysis for a varying number of sires. The offspring number was fixed, at *μ* = 4.63 and 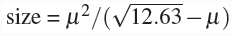 respectively, and the number of sires was varied between 9 and 20. In panel *a.*, the power to reject the null hypothesis of *h*^2^ = 0 is plotted as a function of narrow-sense heritability (*h*^2^), where the true value of *h*^2^>0. In panel *b.*, the power to fail-to-reject the null hypothesis when the true value of *h*^2^ was equal to zero is plotted for varying numbers of sires.

### Power analysis

Power analysis results suggest that increasing the number of sires is the most effective mechanism to increase statistical power. Unfortunately, for heritabilities less then 0.2, very large numbers of sires will be required. The intuition is that substantial amounts of variability between sires is expected just due to sampling alone, and therefore statistical support for a non-zero heritability requires large sample sizes. Despite the lack of statistical power, this approach does have the desirable property of low false positive rates. For example, even with nine sires, we expect to have a nearly 90% chance of failing to reject the null hypothesis on data sets simulated with an among-sire variance equal to zero (Figure 2b). Additionally, if sequencing additional offspring is an option, statistical power can be improved (Figure 3).

**Figure 3.**
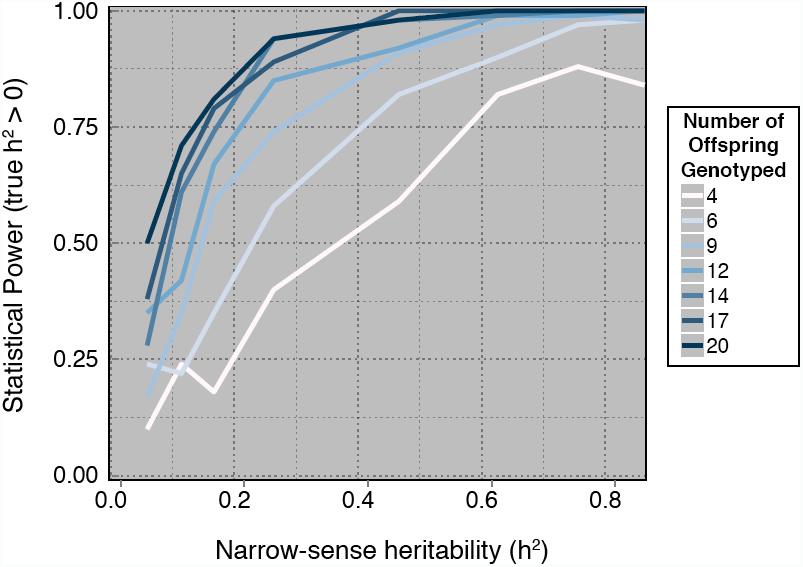
Power analysis for a varying number of offspring. The mean number of offspring genotyped per sire, *μ*, was varied between 4 and 20, while the size parameter for the negative binomial distribution was 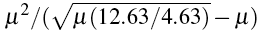. The number of sires was fixed at 9. The power to reject the null hypothesis of *h*^2^ = 0 is plotted as a function of narrow-sense heritability (*h*^2^), where the true value of *h*^2>^0.

## Discussion

In this paper, we present an experimental and statistical methodology for estimating the narrow-sense heritability of binomially distributed traits. We applied this approach to determine whether settlement is a heritable trait in the reef-building coral *O. faveolata*. Although we did not find statistical support for a non-zero, narrow-sense heritability in this trait, a power analysis suggests we lacked a sufficient number of individuals. Our computational method includes code for fitting model parameters, performing model selection using a permutation test, and calculating the expected statistical power for proposed or completed studies. The power calculation method is especially important for studies requiring animal care and use approval and/or those with complex or expensive collection demands.

Previous work suggests that heritable variation exists for a variety of traits across many marine organisms (Foo *et al.*, 2012; Johnson *et al.*, 2010; Kelly *et al.*, 2013; Lobon *et al.*, 2011; McKenzie *et al.*, 2011; Parsons, 1997), including corals (Kenkel *et al.*, 2011; Meyer *et al.*, 2009). Previous studies have found significant heritability for nearly every trait measured in corals (Kenkel *et al.*, 2011; Meyer *et al.*, 2009, 2011; Carlon *et al.*, 2011), but see Csaszar *et al.* (2010). In fact, one study specifically quantified the additive genetic variance in settlement rates of the Pacific reef-building coral *Acropora millepora* and found *h*^2^ = 0.49, however no variance around this mean was estimated (Meyer *et al.*, 2009). It would not be surprising from an evolutionary standpoint if an ecologically important life-history trait such as larval settlement was heritable in other coral species, such as *O. faveoalta*. However, in this study we were unable to detect heritable variation, likely due to insufficient numbers of individuals.

There is a rich quantitative genetics literature on estimating the heritability of binomial traits dating back to Wright (1917) and Fisher (1918); however, the first use of Generalized Linear Models fit to observed presence/absence data is from Gilmour *et al.* (1985). These methods were originally developed for agricultural breeders, where fewer constraints exist on the number of offspring and families used to estimate the heritability–for example the viability of poultry (Robertson & Lerner, 1949), common genetic disorders of Holstein cows (Uribe *et al.*, 1995) and root vigor in sugar beets (Biscarini *et al.*, 2014, 2015). Work by Biscarini *et al.* (2014 and 2015) developed a cross-validation base algorithm for selecting single nucleotide polymorphisms that maximally classified sugar beets into high and low root vigor. Our principle contribution is in terms of model selection, in the form of a permutation test to determine whether statistical support exists for a non-zero narrow-sense heritability, and the methods application to non-model organisms. In such organisms, where breeding, collection, and/or budgetary constraints may exist, such a model-selection procedure is essential.

One caveat is that our methods are somewhat lacking in statistical power. For heritabilities thought to be typical of studies in non-model organisms, well more than 50 individuals may need to be typed across 9 sires, see Figures 2a and 3. However, our methods perform very well with respect to minimizing the type-I error rate, see Figure 2b. Future work should focus on adapting existing methods and developing new methods to allow for smaller sample sizes. This effort is meant to be a project that will grow and develop organically; therefore, we welcome suggestions and contributions and plan regular updates to the statistical methods.

## Acknowledgements

The authors acknowledge funding from the Santa Fe Institute and the Omidyar Group to SVS. Funding was also provided by the National Science Foundation grant DEB-1054766 to MVM, NSF grant DMS-1255187 to JGS, a departmental start-up grant from the Section of Integrative Biology at the University of Texas at Austin to SWD and the PADI Foundation Award to SWD

